# From space to time: Spatial inhomogeneities lead to the emergence of spatio-temporal activity sequences in spiking neuronal networks

**DOI:** 10.1101/428649

**Authors:** Sebastian Spreizer, Ad Aertsen, Arvind Kumar

## Abstract

Spatio-temporal sequences of neuronal activity are observed in many brain regions in a variety of tasks and are thought to form the basis of any meaningful behavior. Mechanisms by which a neuronal network can generate spatio-temporal activity sequences have remained obscure. Existing models are biologically untenable because they require manual embedding of a feedforward network within a random network or supervised learning to train the connectivity of a network to generate sequences. Here, we propose a biologically plausible, generative rule to create spatio-temporal activity sequences in a network model of spiking neurons with distance dependent connectivity. We show that the emergence of spatio-temporal activity sequences requires: (1) individual neurons preferentially project a small fraction of their axons in a specific direction, and (2) the preferential projection direction of neighboring neurons is similar. Thus, an anisotropic but correlated connectivity of neuron groups suffices to generate spatio-temporal activity sequences in an otherwise random neuronal network model.

## Introduction

Ordered sequences of actions are the key to any meaningful behavior. This implies that the task-related neuronal spiking activity in the responsible areas of the brain must also be ordered in temporal activity sequences (Hebb 1949). Indeed, temporal activity sequences have been recorded from different brain regions in various tasks (Hahnloser et al. 2002; Ikegaya et al. 2004; Luczak et al. 2007; Jin et al. 2009; Pastalkova et al. 2008; Harvey et al. 2012; Modi et al. 2014; Bakhurin et al. 2017) (see review by (Bhalla 2017)). Depending on the arousal state of the animal and on the behavioral task, the time scales of the activity sequence may range from few milliseconds to few seconds. The necessity and the ubiquity of the sequential activity patterns in the brain raises the question: what is the origin of such activity sequences in locally random, sparsely connected networks of noisy neurons?

At the simplest, activity sequences of neurons may be attributed to their external inputs. When neurons are tuned to specific properties of an external input, a sequential change in the input can lead to an activity sequence, e.g. temporally ordered firing of place cells in the hippocampus (Dragoi and Tonegawa 2011). However, activity sequences have been observed in several tasks, not involving any specific external stimuli, e.g. in decision making (Jin et al. 2009; Harvey et al. 2012), in learning (Modi et al. 2014), in memory recall (Pastalkova et al. 2008), and in generating bird songs (Hahnloser et al. 2002). These experiments suggest that neuronal networks in the brain are able to generate neuronal activity sequences using some intrinsic mechanisms.

Several computational network models have been proposed to explain the emergence of spontaneous or cue-evoked activity sequences. A feedforward network model (Kumar et al. 2010) is the simplest model that can generate activity sequences, either spontaneously or in response to a short input pulse (Abeles 1991; Diesmann et al. 1999; Kumar et al. 2008). However, given the random and recurrent connectivity in the brain, this architecture is biologically untenable. Recurrent network models with an asymmetric spatial connectivity can exhibit traveling waves (Rinzel et al. 1998; Hutt and Atay 2005; Roxin et al. 2005), which can be considered as a temporal activity sequence. However, in this dynamical regime a network can generate only a single activity sequence, propagating always in the same direction. Recurrent networks tuned to exhibit attractor dynamics (Rabinovich et al. 2006) can generate more diverse temporal activity sequences in response to an external input which steers the spiking activity from one attractor state to another (Zhang 2009). Alternatively, an adaptation in neuronal spiking activity or an activity-dependent short-term-depression of synapses could also be used to create a mechanism to generate activity sequences (Murray and Escola 2017). While such mechanisms provide a natural way to produce both fast and slow sequences, it does not allow for controlling the spatial direction of the activity sequence in a reliable manner. It was shown to be possible to wire the connectivity among neurons representing different attractor states in a way that attractor states switched from one to another in a sequential manner (Murray and Escola 2017). However, it remains unclear how a connectivity necessary for such switching can be achieved. Beyond these networks exhibiting attractor dynamics, more generic echo-state-networks have been trained using a supervised learning algorithm to generate an arbitrary temporal sequence of neuronal activity (Rajan et al. 2016). Thus, at present computational models to generate neuronal activity sequences had to make biologically implausible assumptions as they either required a prewired network connectivity (feedforward network or wiring among different attractor states) or relied on supervised learning (echo-state networks).

Here, we describe a novel mechanism allowing the generation of diverse activity sequences in a recurrent network model without any specific external inputs, any pre-wired networks nor any supervised learning. Specifically, we studied the emergence of activity sequences in a neuronal network with a spatial connectivity profile. We show that when the extent of the spatial connectivity is asymmetric and varying across neurons, spatio-temporal patterns of spiking activity emerge. We identified two conditions that ensured the emergence of spatio-temporal activity sequences (STAS) in such networks of spiking neurons: (1) individual neurons project a small fraction (approximately 2–5%) of their axons in a preferred direction (*ϕ*) and (2) *ϕ*s for neighboring neurons were similar, whereas *ϕ* for neurons further apart were unrelated. These conditions did not depend on the exact composition of neurons in the network model. Both purely inhibitory network models and network models composed of excitatory and inhibitory neurons exhibited STAS, provided the above two conditions were met.

## Results

Can a locally connected random network (LCRN) with excitatory and inhibitory (or only inhibitory) spiking neurons generate STAS in a biologically plausible manner and without embedding feedforward subnetworks (Kumar et al. 2008) or learning such connectivity using a supervised learning algorithm (Rajan et al. 2016)? It is well known that LCRNs can exhibit stable hexagonal patterns of activity bumps (Roxin et al. 2005; Hutt 2008; Spreizer et al. 2017). We hypothesize that such stable spatial activity patterns can be transformed into STAS if the activity bumps could be destabilized. To this end, we investigated the effect of introducing inhomogeneities in the spatial connectivity between neurons on the stability of the activity bumps.

### Spatial distribution of inhomogeneities in neuronal connectivity

We considered an LCRN in which neurons projected a fraction of their axons preferentially in a particular direction (*ϕ*; Figure 1a and Supplementary Figure S2). *ϕ* was chosen from a uniform distribution and assigned to each neuron according to four different configurations (Figure 1b). Random configuration: *ϕ* was randomly and independently assigned to each neuron. Perlin configuration: *ϕ* was assigned to neurons using a gradient noise algorithm such that neighboring neurons had similar values of *ϕ*. Homogeneous configuration: the same *ϕ* was assigned to all neurons. Finally, as a control, we also considered the case in which all neurons projected in all directions with equal probability (Symmetric configuration).

**Figure 1:**
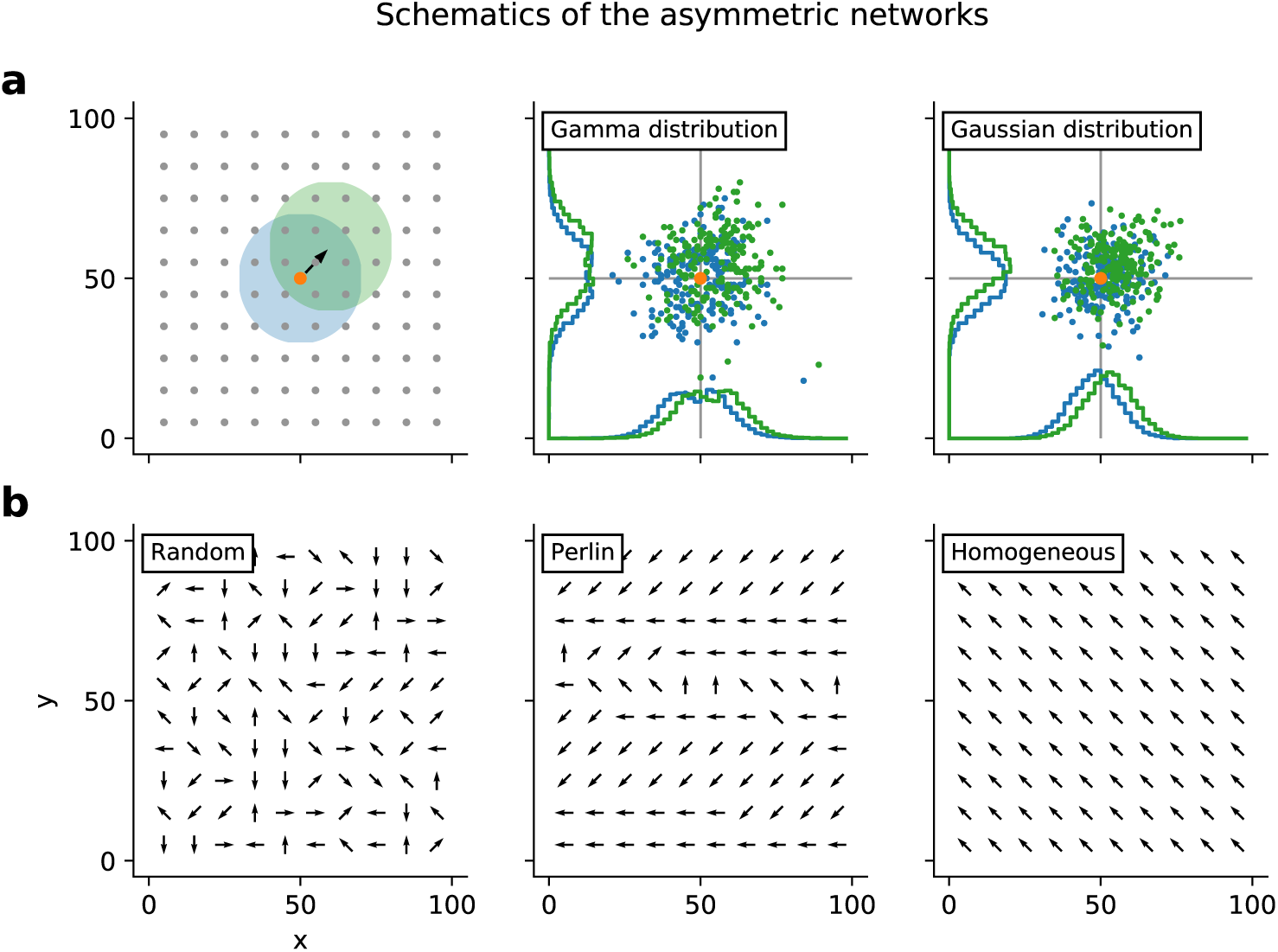
Schematics of the asymmetric network models. **(a:left)** Neurons were arranged on a regular 2-D grid, folded to form a torus. The colored circles indicate the symmetric (blue) and asymmetric (green) spatial connectivity schemes. The pre-synaptic neuron is marked by the orange dot. **(a:center)** Locations of post-synaptic neurons chosen according to the symmetric (green) or asymmetric (blue) connectivity. In this case the distance-dependent connectivity profile varied non-monotonically, according to a Γ distribution. This connectivity profile was used for purely inhibitory network models. **(a:right)** Same as in the center panel, but here the distance-dependent connectivity profile varied monotonically, according to a Gaussian distribution. This connectivity profile was used in the present study for network models with both excitatory and inhibitory neurons. **(b)** Schematic of spatial distribution of connection asymmetries. Each arrow shows the direction in which the neuron makes preferentially most connections (*ϕ*). Here we show examples for random, Perlin and homogeneous configurations.

First, we focused on LCRNs with only inhibitory neurons (I-networks). In these I-networks, we used a connectivity profile which varied non-monotonically with distance, according to a Gamma distribution (Figure 1a:center; see Methods (Spreizer et al. 2017)). After wiring the networks according to each of the four configurations described above, we measured the effective *ϕ* from the spatial distribution of the post-synaptic targets of each neuron. Results are shown in Figs. 2a1-a5. For the random and Perlin configurations, the angle *ϕ* measured from the location of the post-synaptic neurons was uniformly distributed, as was initially specified. For the homogeneous configuration all neurons had identical *ϕ* assigned, but the measured *ϕ* values for individual neurons were slightly different from the assigned value, due to the finite numbers of connections per neuron. For the whole network, *ϕ* was normally distributed around the assigned value, with a very small variance. In the symmetric configuration *ϕ* for the network was uniformly distributed and was different for each neuron, due to the random nature of the connectivity and the finite numbers of connections per neuron.

**Figure 2:**
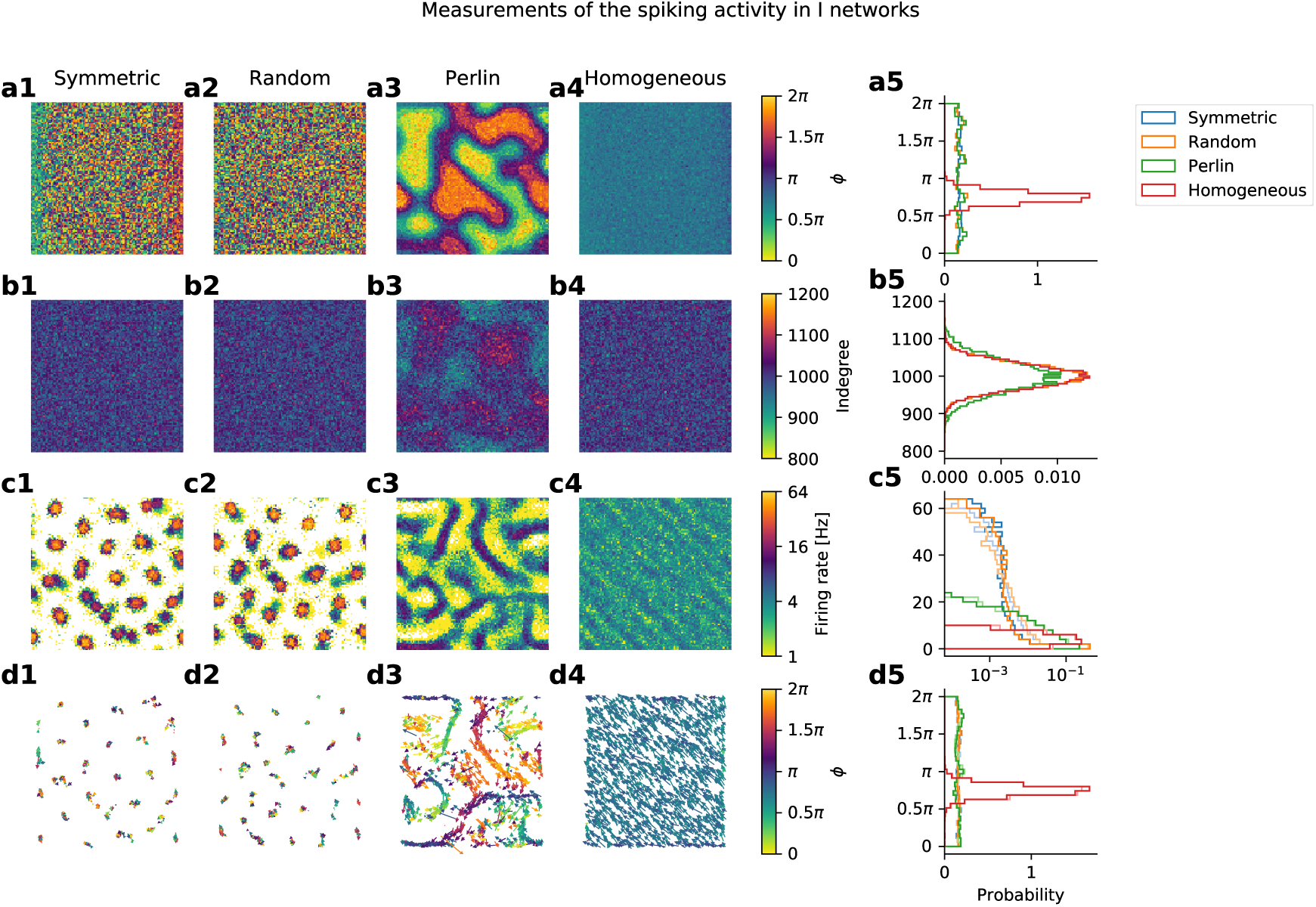
Network structure and spiking activity in I-networks. **(a)** Spatial distribution of connection asymmetries. The square represents the 2-D space of the network. The four panels (**a1-a4**) show the four different configurations of asymmetric connectivity: symmetric, random, Perlin and homogeneous. The panel **a5** shows the distribution of *ϕ*, measured for each neuron from the locations of its post-synaptic neurons. **(b)** Spatial distribution of in-degrees of individual neurons in the four configurations (**b1-b4**). The in-degree distribution was similar for all four configurations (**b5**). Note that in the Perlin configuration, neurons with high and low in-degree were spatially clustered (**b3**). **(c)** Spatial distribution of average firing rates of individual neurons in the four network configurations (**c1-c4**). (**c5**)The distribution of firing rate of all the neurons. **(d1-d4)** Spatially distributed direction of neuronal activity flow in the four configurations. **(d5)** Distribution of the direction of neuronal activity flow independent of space. In symmetric, random and Perlin configurations, activity could move in all possible directions (blue, orange, green), whereas in the homogeneous configuration, activity flowed in a single direction (red). Note that in symmetric and random configurations, despite the presence of all possible directions of projection, the network activity remained locked at certain specific locations (**d1,d2**), unlike in the Perlin configuration, in which a clear and spatially diverse flow of activity emerged (**d3**).

The in-degree distribution was similar across all four configurations (Figs. 2b1-b5). However, in the Perlin configuration, as a consequence of the spatial distribution of *ϕ*, neurons with high and low in-degree distribution were spatially clustered. Thus, the network models were highly similar across all four configurations at the level of neuron properties and their connectivities (same in-degree distribution and fixed out-degree for all neurons).

### Spatial inhomogeneities lead to the emergence of activity sequences

The differences among the four connectivity configurations became evident as we inspected the corresponding network activity dynamics, obtained by activating each neuron in the network with an independent Gaussian white noise (see Methods). In an LCRN with Perlin configuration, time-resolved snapshots of the activity showed transient co-activation of neighboring neurons, referred to as spatial activity bumps (Supplementary Figure S3). Importantly, the spatial bumps were not fixed at a given location, instead as one spatial bump faded, another, similar bump appeared in its immediate vicinity, and so on, thereby creating STAS. Because we did not implement short-term synaptic depression or spike frequency adaptation, the silencing of a spatial bump was a consequence of the network’s dynamical activity state and of the spatial *ϕ*-distribution. Time-averaged firing rates (estimated at over 10 sec) showed that neurons participating in the activity sequences were arranged in stripe like patterns in the network space (Figure 2c), along which the activity sequences flowed.

We used the DBSCAN algorithm (see Methods) to track spatial bumps of spiking activity over time to identify the activity sequences (Figure 3a). In the random configuration for instance we found approximately 23 spatial activity bumps within a time window of 1000 ms (Supplementary Figure S3). The identified activity sequences followed specific paths in the network, visible as stripes in the spatial distribution of average firing rates of individual neurons. Each sequence moved in its own direction, the collection of them forming a uniform distribution of activity sequence movement directions (Figure 2d5).

**Figure 3:**
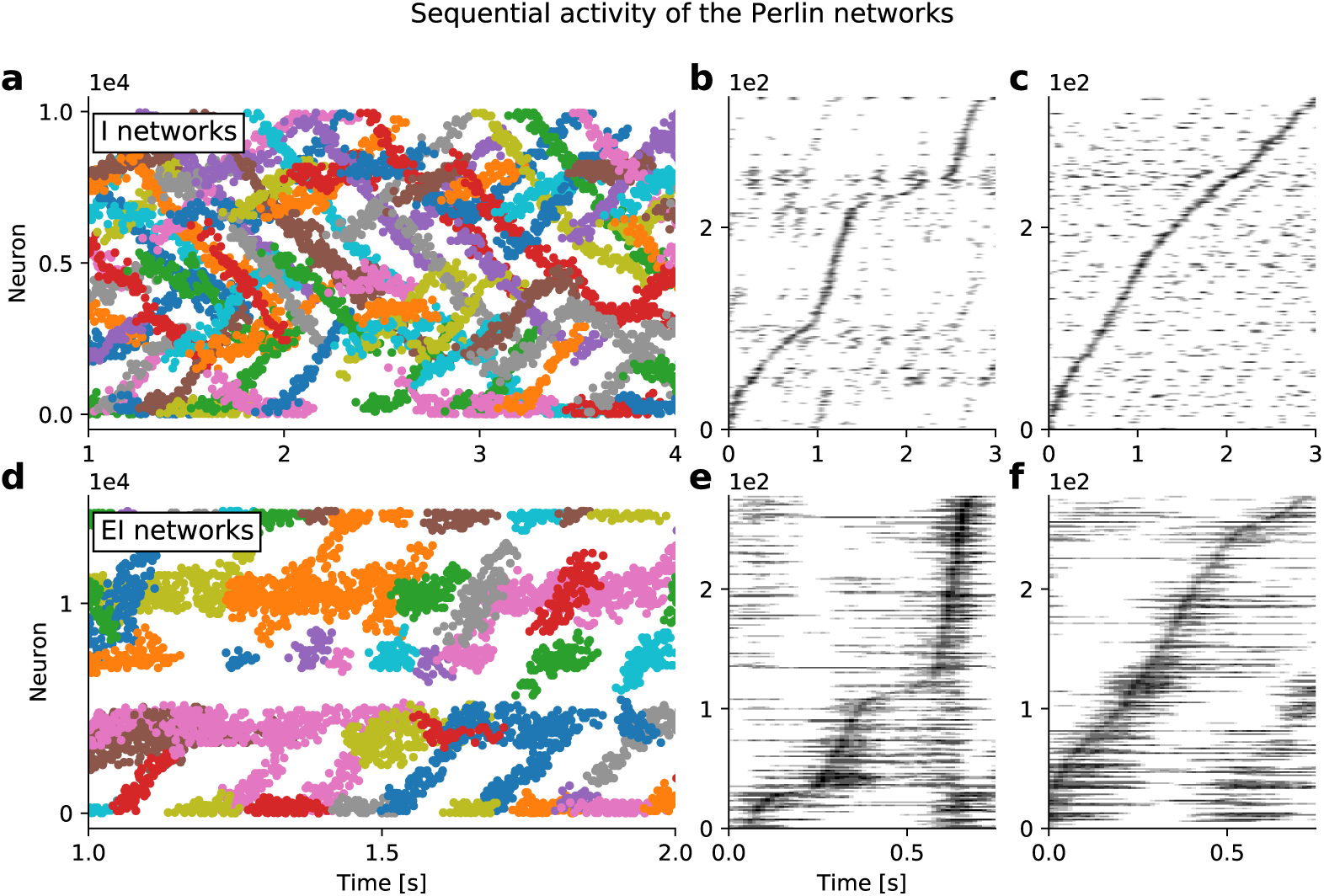
Spatio-temporal sequences of neuronal activity in networks with Perlin configuration. **(a)** Raster plot of spiking activity in an I-network model as a function of time. Each color indicates a cluster of spiking activity in space and time, identified using DBSCAN (see Methods). Note that spikes that were not assigned to any cluster are not shown. **(b)** Activity of approx. 250 neurons confined in a 20 × 20 region. **(c)** Activity of approx. 250 neurons randomly selected from the entire network. **(b, c)** Selected neurons are sorted according to the time of peaked spike counts. **(d, e, f)** Same as in panels **(a, b, c)**, respectively, but here for an EI-network model. Note the shorter time axes in panels **(d, e, f)**, compared to panels **(a, b, c)**, indicating that sequence movement in EI-networks was clearly faster than in I-networks.

In the homogeneous configuration, an extreme case of the connectivity asymmetry, the network activity exhibited multiple moving bumps. Neurons participating in moving activity bumps were arranged in a periodic pattern (Figure 2c4) and the activity sequences flowed along the associated stripes (Figure 2d4). Such patterns of average activity closely resemble with the ’static’ patterns observed in bio-chemical systems (Koch and Meinhardt 1994). Unlike in the Perlin configuration, in the homogeneous configuration all spatial bumps moved in the same direction (Figure 2d5, red trace). Because knowing the movement direction of a single activity sequence was sufficient for knowing the movement directions of all other sequences, the homogeneous configuration effectively exhibited only a single spatio-temporal activity sequence. This type of activity pattern was similar to the traveling waves observed in neural field models with asymmetric connectivity (Roxin et al. 2005; Hutt 2008).

**Figure 4:**
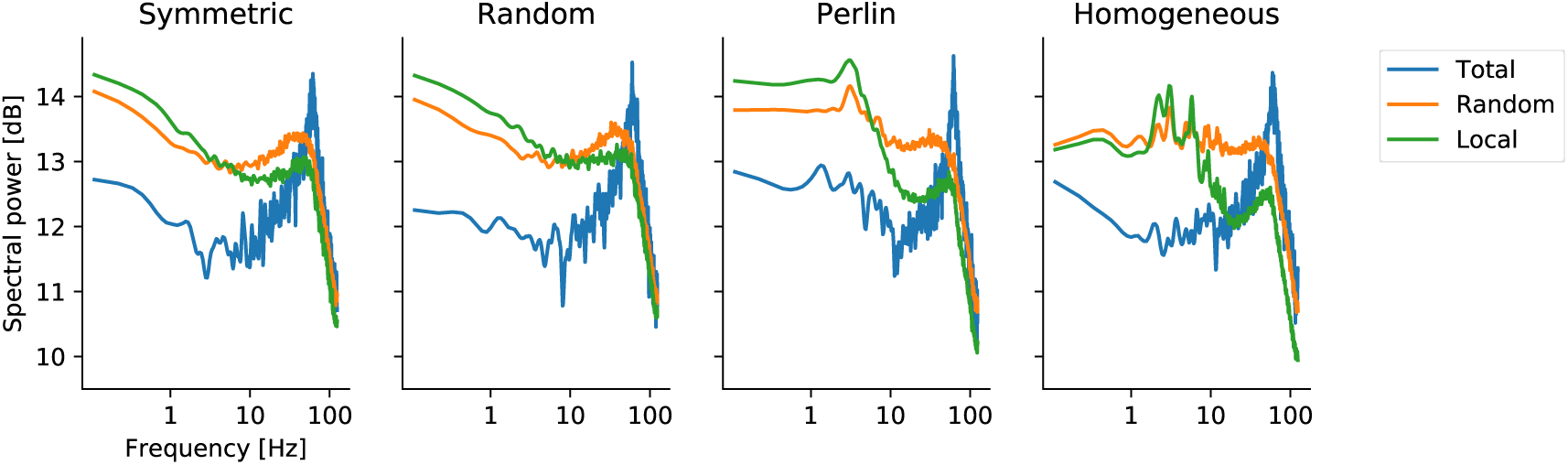
Power spectra of network activity in different spatial inhomogeneity configurations in I-networks. Power spectra of summed spiking activities (bin width = 4ms), with different traces referring to the source of the data: the z-score of the spiking activity of the entire network population (blue trace), of 100 randomly selected neurons from the entire network (orange trace), and of the neurons located in a 10×10 region in the network (green trace). Different traces referred to the scope of data: the z-score of the spike activity of the whole population of neurons (blue trace), of 100 randomly picked neurons (orange trace) and of the neurons located in a 10×10 region in the network (green traces). The spectral power in all network models peaked at approx. 60 Hz (gammaband oscillations). Additionally, in network models with homogeneous and Perlin configurations, an additional, weak low-frequency peak, at around 4–6 Hz, appeared.

When *ϕ* was distributed randomly (random configuration) or when neurons made connections without any directional bias (symmetric configuration), we did not observe any STAS. In both configurations, the network activity was confined to specific neurons, while others were inhibited, giving rise to a long-tailed distribution of average firing rates (Figure 2c5). In both symmetric and random configurations, active neurons were organized in a near hexagonal pattern of spatial activity bumps (Figure 2c1,c2). Such an activity pattern is a consequence of the non-monotonic shape of the effective connectivity (Spreizer et al. 2017). In the random configuration, the spatial organization of the activity bumps was a bit more noisy than in the symmetric configuration. In both configurations, the spatial bumps jittered randomly around a fixed location, resulting in a uniform distribution of bump movement directions (Figure 2d1,d2,d5). Thus, both random and symmetric configurations result in similar types of network activity states.

Similar to the I-network models, an LCRN with both excitatory and inhibitory neurons (EI-networks) also exhibited STAS when excitatory neurons made connections to excitatory neurons preferentially in one direction and *ϕ*-values were distributed according to the Perlin configuration (Figure 3d). In both EI- and I-network models, the activity sequence could be extracted from only a few neurons chosen from a small neighborhood (Figure 3b,e) or randomly from the whole network (Figure 3c,f). When the spiking activity was sampled from the entire network and neurons were ordered according to their peak firing rates (as is often done with experimental data (Harvey et al. 2012; Bakhurin et al. 2017)), the velocity of the activity sequence appeared to be quite constant (see Figure 3c,f). Experimental data suggest that the velocity of temporal sequences can vary over time (Bakhurin et al. 2017). In our network model, we also found that when about 250 active neurons were sampled randomly from a small network neighborhood, the velocity of the activity sequences varied as a function of time (Figure 3b). However, this varying velocity could be an artefact of the finite size effect and of the non-uniform sampling of the sequences (see Figure 3b,c). In general, the activity sequences in EI-networks model were faster than those in I-networks, because the activity sequences in EI-networks relied on recurrent excitation, whereas in I-networks they relied on the lack of the recurrent inhibition (in our I-networks neuronal connectivity varied non-monotonically with distance, according to a Gamma distribution, therefore there was a small connection probability among neighboring neurons).

### Conditions for the emergence of spatio-temporal activity sequences

These results suggested that the emergence of STAS in LCRNs required two conditions to be met: (1) each neuron projects a fraction of its axons preferentially in a specific direction (*ϕ*) and (2) neighboring neurons preferentially project in similar directions. These two conditions imply a spatially correlated anisotropy in the projection patterns of neurons in the network. Indeed, upon systematic variation of a wide range of input parameters and excitation-inhibition balance, we found that, as long as these two conditions were met, irrespective of the composition of neurons in the LCRN, STAS invariably emerged (Supplementary Figure S4).

### Co-existence of activity sequences and network oscillations

The rasters of spiking activity in both I-networks and EI-networks indicated the presence of slow oscillations in Perlin (Figure 3) and homogeneous (not shown) configurations. Therefore, we measured the spectrum of the summed network activity. The network activity was obtained by different procedures: by summing the activity of all neurons (Figure 4, blue trace), by summing the activity of the neurons from a 10 × 10 region in the network (Figure 4, green trace), and by summing the activity of 100 randomly chosen neurons from the entire network (Figure 4, orange trace). For both I-networks (Figure 4) and EI-networks (Supp Figure S5), neuronal population activity in all four configurations exhibited clear oscillations in the gamma frequency band (30–60 Hz). These oscillations were a global property of the network, as partial sampling of the neurons showed only weak signs of oscillatory activity. (Figure 4, orange and green traces). Moreover, in homogeneous and Perlin configurations, signs of low-frequency oscillations at around 4–6 Hz were observable. These were presumably a consequence of the periodic boundary conditions, i.e. the period of slow oscillations was determined by the sequence propagation velocity (see below) and the spatial network scale. These results suggest that both STAS and global oscillations can co-exist in the same network model, however, one did not automatically imply the other.

### Asymmetry in connectivity determines the velocity of spatiotemporal sequences

Next we investigated how the amount of shift in the connectivity affects the STAS. To this end we shifted the connectivity extent by 1 and 2 grid points. When there was no shift in the connectivity, network did not exhibit any sequential activity and the activity bumps jittered around a fixed value with a small velocity (Figure 5a,b blue). However, shifting the connectivity by 1 grid point was sufficient to induce sequential activity in both homogeneous and Perlin configurations. The velocity of STAS was higher in homogeneous configuration than in Perlin configuration (Figure 5a,b orange). When we increased the shift in connectivity by 2 grid points, the mean and the variance of velocity increased in both Perlin and homogeneous configurations (Figure 5a,b green). These results suggest that the degree of asymmetry in the connectivity controls the velocity of STAS.

**Figure 5:**
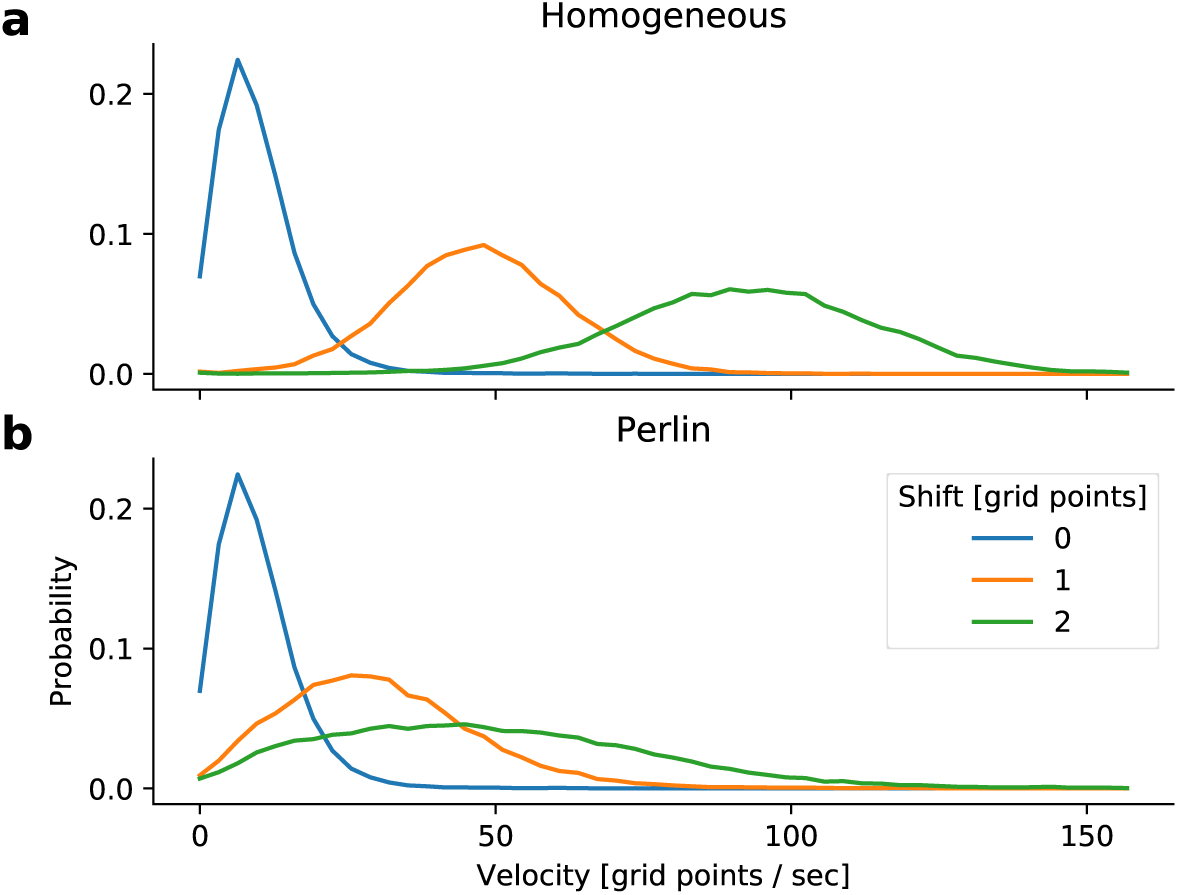
Velocity of neuronal activity sequences. Distribution of the velocity of neuronal activity sequences in I-networks with homogeneous (**a**) and Perlin (**b**) configurations. The velocity of the activity sequences increased with the increase in connectivity asymmetry. Note that the velocity of activity bump movements in networks with symmetric connections (blue traces) were identical in networks with homogeneous and Perlin configurations. However, for any non-zero degree of asymmetry (orange and green traces) the velocity of activity bump movements was higher in networks with the homogeneous configuration.

### Effect of spatial correlation in connection asymmetry on spatio-temporal activity sequences

Next, we determined how the spatial correlations in *ϕ* affect the number and velocity of STAS. To this end we systematically varied the spatial scale of the Perlin noise (see Methods). This enabled us to systematically go from a random configuration to a homogeneous configuration (Figure 6a top). To count the number of STAS we rendered the activity in a 3-dimensional space (two space dimensions and one time dimensions) and used DBSCAN algorithm to calculate clusters (which are the STAS) in this 3-D space. We found that the number of STAS and their velocity decreased monotonically as we reduced Perlin scale (Figure 6b,c). This decrease in number and velocity of STAS occurred because reduction in the Perlin scale reduced the number of neighboring neurons with similar *ϕ*. This, in turn, reduced the velocity of the STAS (Figure 6b), because the input in the direction specified by *ϕ* decreased and the postsynaptic neurons had to integrate over a longer time to elicit response spikes. Moreover, because of fewer inputs in the direction *ϕ* many putative sequences showed weak spiking activity which could not be classified as a distinct spatio-temporal sequence. Furthermore, reduction in Perlin scale also increased the variance of movement directions (Figure 6c). These results show, first, that even a small scale spatial correlation in *ϕ* suffices to induce STAS but, second, if the spatial correlation scale is too small, such sequences may not move quickly enough to be noticed as sequences. For functionally relevant STAS, the spatial correlation in *ϕ* should be about 20 which is about 1/6th of the network size.

**Figure 6:**
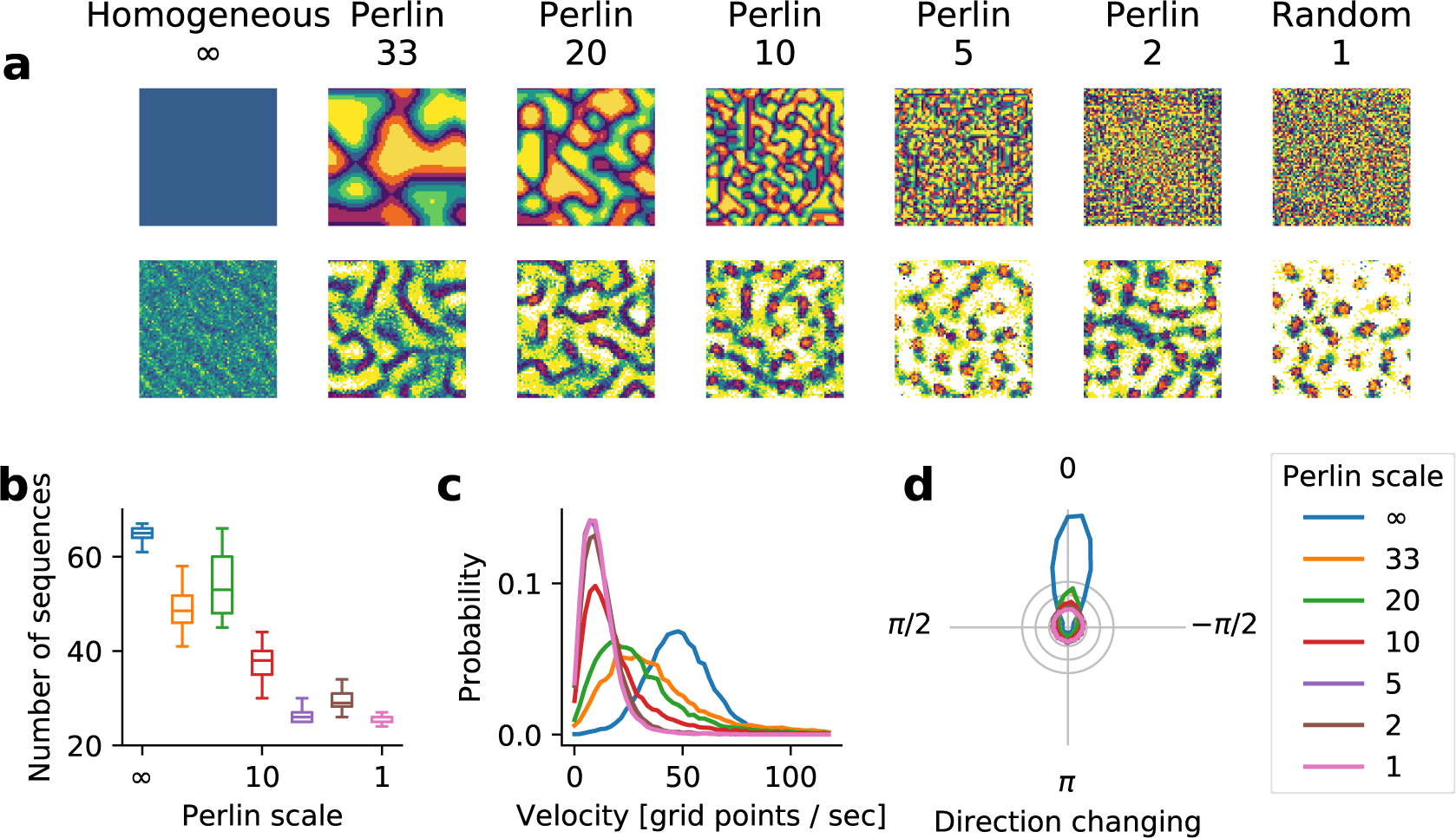
Effect of spatial scale of correlations in *ϕ* on the emergence and velocity of STAS in I-networks. **(a)** The top row shows the spatial distribution of *ϕ* for different scales of Perlin noise. The Perlin scales decreased from left to right as reflected in the size of single color blobs. The Perlin scale is indicated in terms of grid points in the network. The bottom row shows the spatial distribution of average firing rates in each of the seven configurations. **(b)** The number of STAS observed in 1 sec. for different Perlin scales. The box plot shows that statistics of STAS estimated over 90 epoch of 1 sec. each. Different colors indicate the scale of the corresponding Perlin noise. **(c)** The distribution of the velocity of STAS. Different colors indicate the scale of the corresponding Perlin noise. **(d)** The distribution of STAS directions in polar plots. In a homogeneous configuration, most sequences moved in a single direction (blue curve). As the Perlin scale decreased, the distribution of movement direction became more widely distributed, indicating an increase in the number of sequences that moved in different directions.

### Why does spatially correlated connection asymmetry give rise to spatio-temporal activity sequences?

To get more insights into the mechanisms underlying the emergence of the STAS in Perlin and homogeneous networks we estimated the eigenvalue spectrum of the network’s connectivity matrix. For an Erdós-Renyi type random network, eigenvalues of the connectivity matrix are distributed in a circle (Rajan and Abbott 2006). In an inhibition dominated network, extra inhibition introduces very large negative eigenvalues that contribute to the stability of the network activity dynamics (Pernice et al. 2011). Here, we found that for an LCRN without any directional connectivity (symmetric configuration), most of the eigenvalues were confined within a circle, but the local nature of the connectivity introduced several eigenvalues outside the circle, with large real parts and small imaginary parts. As we introduced the spatial asymmetry into the connectivity, the imaginary component of large eigenvalues (those outside the circle) increased (Figure 7a). The emergence of large complex eigenvalues outside the main circle is indicative of meta-stable dynamics (Rajan and Abbott 2006). Moreover, the number of large eigenvalues outside the main lobe (circle in this case) of the eigenvalue spectrum is equal to the number of ’communities’ of neurons in a network (Zhang et al. 2014). This suggests that in both Perlin and homogeneous configurations correlations in the spatial distribution of *ϕ*s created many communities (neuronal assemblies), the dynamics of which are meta-stable.

**Figure 7:**
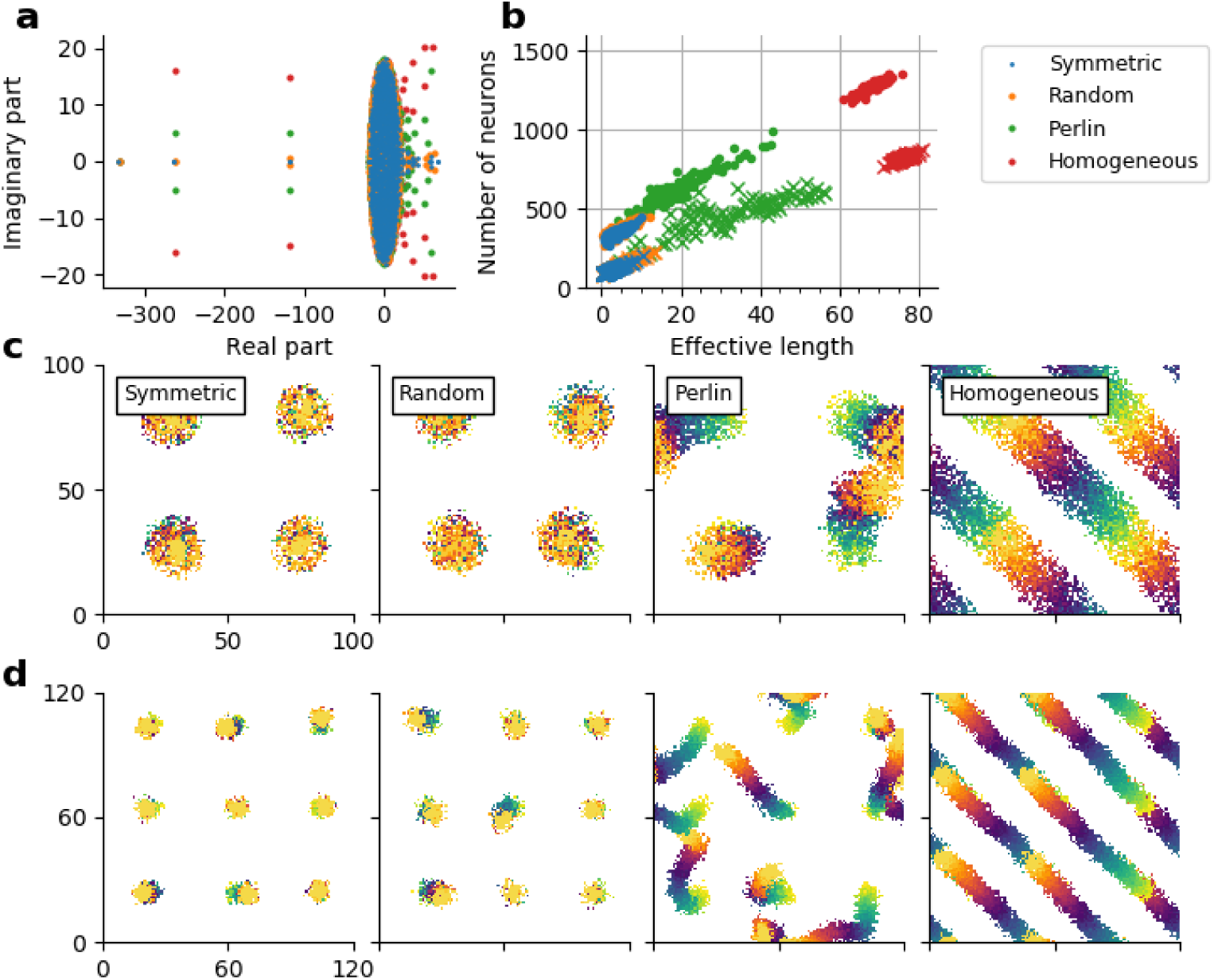
Spatial clustering of *ϕ* results in feedforward pathways in otherwise locally connected random networks. **(a)** The eigenvalue spectrum of the connectivity matrix of 1000 inhibitory neurons randomly taken from symmetric (blue dots), random (orange dots), Perlin (green dots) and homogeneous (red dots) I-networks. **(b)** Number of unique target neurons participating in a feedforward path (y-axis) as a function of the effective length of the feedforward path (Euclidean distance between the centroids of *F*_1_ and *F*_50_ (see Methods). Feedforward path in I-network (dots), feedforward path in El-network (crosses). The four colors indicate the network configurations. Note that distinctly more unique neurons with longer path length of the sequential activity movement were observed in Perlin and homogeneous configurations. **(c)** Effective feedforward pathways in an l-network model with the four configuration (see Methods). Feedforward paths starting from four different locations are shown. The starting neuron set *F*_1_ is shown in yellow, the final set *F*_50_ is shown in orange. Effective feedforward pathways were visible as trails changing color from yellow to orange. The starting neuron set *F*_1_ consisted of 64 neurons located in 8 × 8 region of the network. **(d)** Same as in **c**, but for an El-network model with nine different starting set locations

Given the large size of our network models, it is computationally highly demanding to test this hypothesis by measuring all eigenvalues of an LCRN, identifying and counting the neuronal assemblies and determining the effective feedforward networks associated with their STAS. To simplify the problem, we estimated the probability of finding such a feedforward network *pFF* in our I- and EI-network models. To this end, we used an iterative procedure to find feedforward networks in our network models, the details of which are described in the Methods section. Briefly, we started with a set of 64 neurons (*F_i_*) located in a small, 8 × 8 region in the network. Then we identified a set of all post-synaptic neurons (*P_i_*) receiving input from any of the neurons in the first set *F_i_*. From the set *P_i_* we selected the 64 neurons (*F_i_*_+1_) that were most frequently connected to the neurons in the input *F_i_* (see Methods for details). We repeated this procedure 50 times, starting at 100 randomly selected different locations of the initial 8 × 8 regions (see Methods, Figure 7b,c). Thus, we identified feedforward networks with excitatory connections from *F_n_* to *F_n+1_* in EI-networks and feedforward networks with inhibitory connections from *F_n_* to *F_n_*_+1_ in l-networks.

In the homogeneous configuration we always found a feedforward path capable to creating a STAS (Effective length > 16; see Methods and Figure 7c-d). Indeed, in the homogeneous configuration, the probability to find a feedforward path: *pFF* was 1.0 (for both EI- and I-networks). Moreover, these feedforward paths were very long (Figure 7b, red dots and crosses). In the Perlin configuration, there were fewer (*pFF* ≈ 0.8 EI-networks; *pFF* ≈ 0.66 I-networks) and shorter (Figure 7b, green dots and crosses) feedforward paths, but they pointed in different directions (Figure 7b-d). By contrast, in both symmetric and random configurations, no feedforward pathways were observed (*pFF* = 0 for both EI- and I-networks). In these latter two configurations, neurons participating in *F*_1_ to *F*_50_ were confined to a small space (indicated by the overlap of the color blobs in Figure 7c,d). Ultimately, it was the existence (or non-existence) of these feedforward pathways that determined the properties of the STAS in the four different configurations.

In EI-networks, within the feedforward path excitatory neurons from *F_n_* projected to excitatory neurons within *F_n_*_+1_ with a higher probability than outside *F_n_*_+1_, thereby creating a path of high excitation between adjacent groups. When an external input was strong enough STAS was observed along such paths of high excitation. By contrast, in I-networks within the feedforward path, inhibitory neurons from *F_n_* projected to inhibitory neurons within *F_n_*_+1_ with a higher probability than outside *F_n_*_+1_, thereby, creating a path of high inhibition from *F*_1_ to *F*_50_. Because the out-degree of neurons was fixed, the concentration of inhibitory connections within the path from *F*_1_ to *F*_50_ created a region of low recurrent inhibition in the vicinity of the path, along which inhibitory STAS emerged. Thus, the abundance of feedforward paths in networks with Perlin and homogeneous configurations provided a structural substrate for the emergence of the rich repertoire of STAS.

Feedforward networks are simple but powerful computing devices (Abeles 1991; Kumar et al. 2010). Moreover, such feedforward networks are also thought to be the structural substrates of the *phase sequences* that Hebb proposed to ’neuralize this behavior’ (Hebb 1949). Therefore, there is a general interest in understanding how feedforward networks may emerge in an otherwise randomly connected networks. To this end, a number of computational studies have investigated whether Hebbian synaptic plasticity can generate such feedforward networks. These attempts are usually successful in creating feedforward networks in small random recurrent network (Masuda and Kori 2007; Clopath et al. 2010; Liu and Buonomano 2009; Fiete et al. 2010) but do not scale up for large recurrent networks (Kunkel et al. 2011). Our observations of feedforward networks in an LCRN with Perlin configuration provides a much simpler generative mechanism that can create feedforward networks in large random neuronal networks without considering any synaptic plasticity.

## Discussion

Here we have shown that spatial inhomogeneities in network connectivity can lead to the emergence of STAS in the network. Unlike existing models of sequence generation, which require either manual wiring of neurons or supervised learning, we provide a simple generative rule that renders an LCRN with the ability to generate STAS. We showed that when (1) individual neurons project a small fraction (approximate 2–5%) of their axons in a preferred direction, *ϕ* (i.e. the connectivity is asymmetric), and (2) *ϕ*s of neighboring neurons are similar, whereas *ϕ* of neurons further apart were unrelated (i.e. the network is anisotropic), the network will generate STAS. That is, asymmetric but spatially correlated connectivity of neurons translates into sequential spiking activity. Note that, the spatial asymmetry of neuronal connectivity can also be achieved when a neuron makes stronger instead of more synapses in the preferred direction (*ϕ*). Under this mechanism, the number of STAS and their propagation velocity are determined by the extent of the connectivity asymmetry and the spatial scale of the *ϕ*-correlations (Figure 6).

Our proposed sequence generation mechanism demands another mechanism which enables a group of neurons to make more or stronger synapses in a common direction *ϕ* in the first place. Axonal and dendritic arbors of neurons are almost never symmetric in space (Mohan et al. 2015) and dendritic arbors of some prominent neuron types are highly similar. However, it is not possible to infer from the available data whether neighboring neurons have similar orientation of axons. Experimental data suggest that neurons born together tend to share their inputs (Li et al. 2012). In addition, activity dependent plasticity may also lead to the formation of a few stronger synapses, possibly (but not necessarily) associated with a preferred projection direction. However, such mechanisms, even if viable, will only hardwire one specific set of STAS. In the following, we propose a more general and, more importantly, dynamic mechanism that may lead to asymmetric and spatially correlated connectivity of neurons that not only generates STAS, but may rapidly switch from one set of sequences to another. Consider a network in which neurons have symmetric dendritic and axonal arbors (Figure 8a). Such a network would not support activity sequences (Figure 2) and stimulus evoked activity will be confined to the stimulated region of the network. In this network, the release of a neuromodulator (e.g. dopamine or acetylcholine) will create a phasic increase in the neuromodulator levels in small patches (Figure 8b, yellow blobs). Such patches naturally arise because of the non-uniform distribution of axons releasing the neuromodulator and the diffusion in the neural tissue which presents an inhomogeneous medium. A similar patchy spatial profile of dopamine has been recently observed experimentally *in vivo* (Patriarchi et al. 2018). Most synapses within the regions of high neuromodulator concentration will be potentiated (schematically shown in Figure 8b, blue neurons) and, hence, create an asymmetric, spatially correlated connectivity for as long as the neuromodulator concentration remains high. That is, along the spatial gradient of the neuromodulator concentration (Figure 8b, arrows), effective connectivity may be modified to generate STAS.

**Figure 8:**
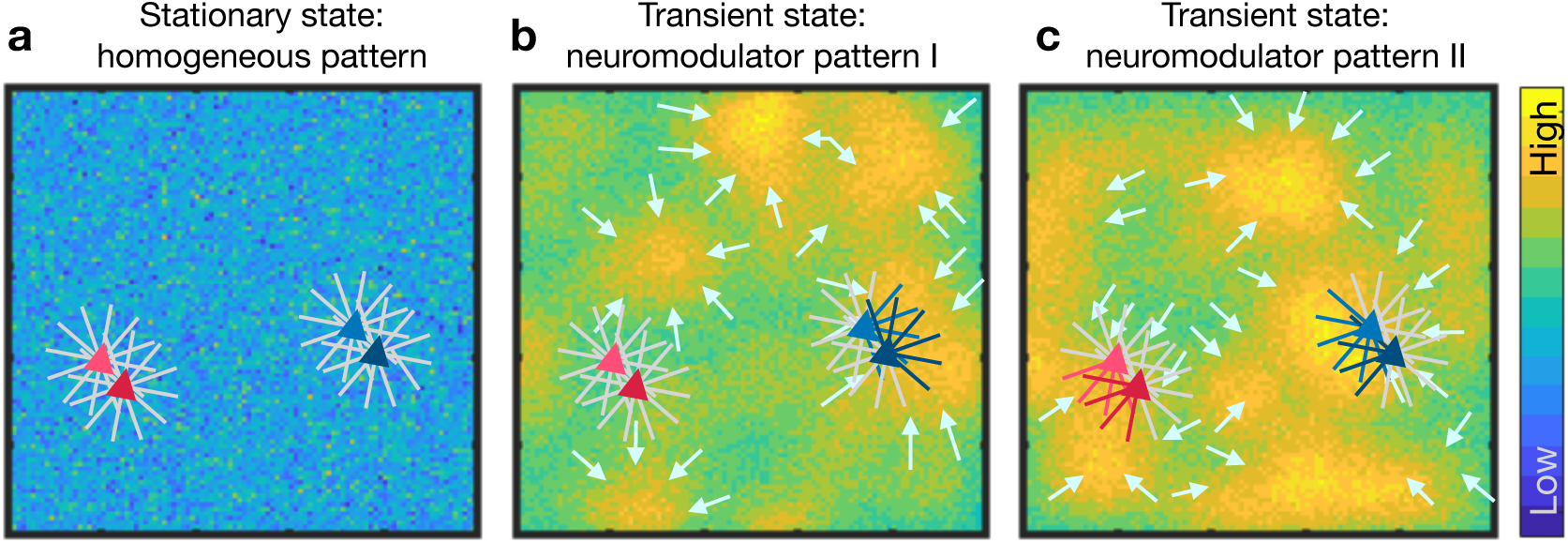
Dynamic reorganization of activity sequences in a recurrent network. **(a)** Schematic of a network in which neurons connect in all direction equally. The blue background shows the baseline level of a certain neuromodulator substance. Two pairs of neurons (blue and red triangles) are shown, the axons of which project in all directions uniformly. This is equivalent to the symmetric configuration and, hence, no sequential activity will emerge. **(b)** Non-uniform distribution of concentration of the neuromodulator in different parts of the network, as indicated by the colormap. The colored lines indicate the enhanced synaptic strength in specific directions. Asymmetric connectivity of neighboring neurons caused by such non-uniform neuromodulator concentration distribution may result in activity sequence in some regions of the network (e.g. neurons marked in blue). The short arrows mark the potential flow of a neuronal activity sequence along the spatial gradient of the neuromodulator concentration. **(c)** Same as in **b** for a different pattern of neuromodulator concentration which may lead to a different flow of neuronal activity, resulting in the appearance of activity sequences in a new set of neurons (e.g. those marked in red) and a change in direction of the sequence in others (e.g. those marked in blue).

This neuromodulator based mechanism to generate spatially correlated asymmetric, anisotropic connectivity automatically provides a mechanism to rapidly switch between sequences. A change in the spatial profile of the neuromodulator concentration will potentiate another set of synapses, possibly leading to the recruitment of new neurons in the activity sequence (Figure 8b,c red neurons), or to the assignment of neurons to a different sequence (Figure 8b,c, blue neurons). In the discussion above, we assumed that the neuromodulator enhances synaptic strengths, but the same argument holds also when the neuromodulator suppresses synaptic strengths. Thus, neuromodulators may play a crucial role, not only in the formation of STAS, but also in rapidly switching between different sets of sequences. Moreover, by their spatial concentration distribution, neuromodulators can also control the speed of the activity sequence. This idea is consistent with experimental observations, e.g. the finding that acetylcholine is important for retinal waves (Ackman et al. 2012).

The key prediction of our network model is that in brain regions generating STAS, neurons should have asymmetric but spatially correlated network connectivity. Such correlated asymmetry can be observed in at least two different forms: (1) asymmetric but similar axonal or dendritic arbors of neighboring neurons, and (2) neighboring neurons receiving strong synapses from common sources and sending out strong synapses to common targets. Secondly, we predict that neuromodulators play a crucial role in the generation and control of STAS in an otherwise isotropic networks. This can be tested by experimentally controlling the spatial profile of the corresponding neuromodulator release pattern by optogenetic stimulation of its source neurons.

In summary, we propose a simple generative rule that enables neuronal networks to generate STAS. How these spontaneously generated sequences interact with stimuli and how we can create stimulus - sequence associations is an interesting and involved question that will be addressed in future work. Similarly, more work is needed to determine the role of neuronal and synaptic weight heterogeneities in shaping spontaneous and stimulus-evoked neuronal activity sequences, either with or without changes in neuromodulator concentration distributions.

## Methods

### Neuron model

Neurons in the recurrent networks were modelled as ‘leaky-integrate-and-fire’ (LIF) neurons. The sub-threshold membrane potential (*v*) dynamics of LIF neurons are given by:

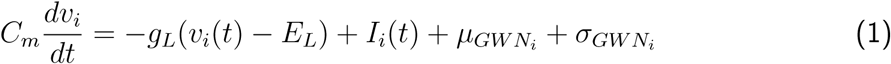

where 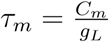 denotes the membrane time constant, *C_m_* the membrane capacitance, *g_L_* the leak conductance, *E_L_* the leak reversal potential, and *I*(*t*) the total synaptic current. The neuron parameters are listed in Table 1, top.

**Table 1:**
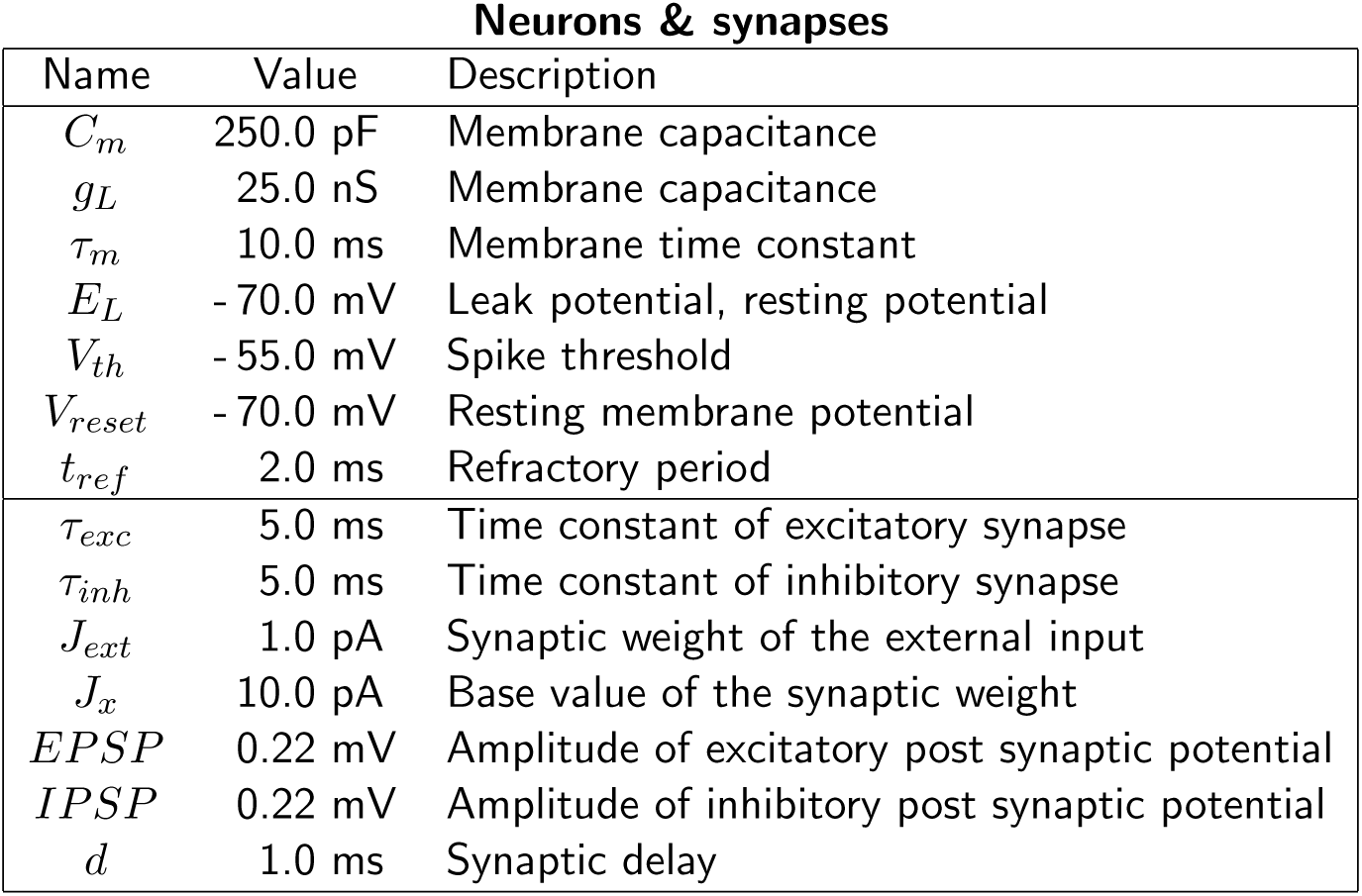
Parameter values for the neurons (top) and for the synapses (bottom) in both network models.

| Neurons & synapses |  |  |
| --- | --- | --- |
| Name | Value | Description |
| $C_m$ | 250.0 pF | Membrane capacitance |
| $g_L$ | 25.0 nS | Membrane capacitance |
| $\tau_m$ | 10.0 ms | Membrane time constant |
| $E_L$ | -70.0 mV | Leak potential, resting potential |
| $V_{th}$ | -55.0 mV | Spike threshold |
| $V_{reset}$ | -70.0 mV | Resting membrane potential |
| $t_{ref}$ | 2.0 ms | Refractory period |
| $\tau_{exc}$ | 5.0 ms | Time constant of excitatory synapse |
| $\tau_{inh}$ | 5.0 ms | Time constant of inhibitory synapse |
| $J_{ext}$ | 1.0 pA | Synaptic weight of the external input |
| $J_x$ | 10.0 pA | Base value of the synaptic weight |
| $EPSP$ | 0.22 mV | Amplitude of excitatory post synaptic potential |
| $IPSP$ | 0.22 mV | Amplitude of inhibitory post synaptic potential |
| $d$ | 1.0 ms | Synaptic delay |

Synapses in the network were modeled as current transients. The temporal profile of the current transient in response to a single pre-synaptic spike was modeled as an *α* function. We adjusted the synaptic currents to obtain weak synapses such that both a unitary inhibitory post-synaptic potential (IPSP) and a unitary excitatory postsynaptic potential (EPSP) had an amplitude of 0.22 mV. The synapse parameters (synaptic strength, time constant and delay) were fixed throughout the simulations and are listed in Table 1, bottom.

### Network architecture

We studied two types of recurrent network models in which the connection probability between any two neurons depended on the physical distance between them. Neurons in both network models were placed on a regular square grid. To avoid boundary effect, the grid was folded to form a torus (Kumar et al. 2008). In both network types, multiple connections were permitted (Supplementary Figure S1), but self-connections were excluded.

#### Networks with only inhibitory neurons

The first type network model (I-network) was composed of only inhibitory neurons. These neurons were arranged on a 100×100 grid (*npop* = 10,000). Each neuron projected to 1,000 other neurons (corresponding to an average connection probability in the network of 10%). The distance-dependent connection probability varied according to a Γ distribution (Spreizer et al. 2017) with the following parameters: *κ* = 4 for the shape and *θ* = 3 for the spatial scale. All I-network parameters are summarized in Table 2.

**Table 2:** Parameter values for the networks (top), for the connections (middle) and for an external input (bottom) in I network model.

| I network model |  |  |
| --- | --- | --- |
| Name | Value | Description |
| Neuron model |  | Integrate and fire |
| $nrow, ncol$ | 100 | number of rows/columns in a network layer |
| $npop$ | $ncol \times nrow = 10,000$ | number of neurons |
| Synapse model | | $\alpha$ function, current-based model |
| $\kappa$ | 4 | Shape for gamma distribution function |
| $\theta$ | 3 | Scale for gamma distribution function |
| $p_{conn}$ | 0.1 | Average connection probability of target neurons |
| $n_{conn}$ | $p_{conn} \times npop = 1000$ | Number of recurrent connections per neuron |
| $J_{rec}$ | $-J_x = -0.22$ mV | Synaptic weight of recurrent inhibitory connections |
| $\mu_{GWN}$ | 700.0 pA | Mean of external GWN input |
| $\sigma_{GWN}$ | 100.0 pA | Standard deviation of external GWN input |

#### Networks with both excitatory and inhibitory neurons

The second type network model (EI-network) was composed of both excitatory and inhibitory neurons. Excitatory and inhibitory neurons were arranged on a 120×120 (*npop_E_* = 14,400) and on a 60×60 grid (*npop_I_* = 3,600), respectively. Each neuron of the excitatory and inhibitory populations projected to 720 excitatory and 180 inhibitory neurons (average connection probability 5%). The connection probability varied with distance between neurons according to a Gaussian distribution (Kumar et al. 2008; Schnepel et al. 2015; Spreizer et al. 2017). The space constant (standard deviation of the Gaussian distribution) for excitatory targets was *σ_E_* = 12 and for inhibitory targets *σ_I_* = 9. We considered a high probability of connections within a small neighborhood, therefore, these networks were referred to as locally connected random networks (LCRN (Mehring et al. 2003)). All EI-network parameters are summarized in Table 3. Whenever possible, we used parameters corresponding to a standard EI-network (Brunel 2000).

**Table 3:** Parameter values for the networks (top), for the connections (middle) and for an external input (bottom) in EI-network model.

| EI network model |  |  |
| --- | --- | --- |
| Name | Value | Description |
| Neuron model |  | Integrate and fire |
| $nrow_E, ncol_E$ | 120 | number of rows/columns in exc. network layer |
| $nrow_I, ncol_I$ | 60 | number of rows/columns in inh. network layer |
| $npop_E$ | $ncol_E \times nrow_E = 14,400$ | number of excitatory neurons |
| $npop_I$ | $ncol_I \times nrow_I = 3,600$ | number of inhibitory neurons |
| $npopratio$ | $npop_E : npop_I = 4 : 1$ | Ratio of exc. - inh. neurons |
| Synapse model | | $\alpha$ function, current-based model |
| $\sigma_E$ | 12 | Space constant for excitatory targets |
| $\sigma_I$ | 9 | Space constant for inhibitory targets |
| $p_{conn}$ | 0.05 | Connection probability of target neurons |
| $n_{connE}$ | $p_{conn} \times npop_E = 720$ | Connection number of excitatory targets |
| $n_{connI}$ | $p_{conn} \times npop_I = 180$ | Connection number of inhibitory targets |
| $g$ | 6 | Ratio of recurrent inhibition and excitation |
| $J_E$ | $J_x = 0.22$ mV | Synaptic weights of excitatory targets |
| $J_I$ | $g \times J_E = -1.32$ mV | Synaptic weights of inhibitory targets |
| $\mu_{GWN}$ | 350.0 pA | Mean of external GWN input |
| $\sigma_{GWN}$ | 50.0 pA | Standard deviation of external GWN input |

### Asymmetry in spatial connections

Typically, in network models with distance-dependent connectivity, the connection probability is considered to be isotropic in all directions. In the network model studied here, however, we deviated from this assumption and introduced spatial inhomogeneities in the recurrent connections. Specifically, we considered a scenario in which the neuronal connectivity was asymmetric in the sense that each neuron projected a small fraction of its axons in a particular direction *ϕ* (Figure 1a,b). At the same time we ensured that the out-degree of each neuron was the same as in an LCRN with isotropic connectivity. The fraction of extra connections in the direction *ϕ* depended on the shift in the region of post-synaptic neurons (green circle in Figure 1a). To quantify the change in connection probability, we estimated the average distance-dependent connectivity in the symmetric configuration (*S*) and in the homogeneous configuration (*H*). The change in connectivity was then measured as (*S − H*)/*S*.

Shifting the connectivity region of a neuron by one grid location in our network model means that the probability of a neuron to make a connection in that direction was increased or decreased by some amount Δ*p* = 0 − 100%, depending on the distance between the neurons. At short distances, the connection probability almost doubled, whereas at distances between 10–20 grid points, there was only a very small change in the connection probability (Supplementary Figure S2). At larger distances (> 20 grid points), the connection probability changed by a large amount (Supplementary Figure S2). This is because at such distances the connectivity is sparse (connection probability < 0.01) and the measure (*S − H*)/*S* amplifies small changes for small *S*. Because we maintained the out-degree of the neurons, an increase in the connection probability in one direction implied a reduction in connection probability by the same amount in the opposite direction.

Note that an increased connection probability also increased the probability to form multiple connections in the close neighborhood of the projecting neuron (Supplementary Figure S1). The preferred direction (*ϕ*) for each neuron was chosen at random from a set of eight different directions, considering that neurons were positioned in a grid pattern. All other synaptic parameters, such as the number of total connections, the space constant of the connectivity kernel and the synaptic weights were identical for all neurons.

### Spatial distribution of asymmetry in spatial connections

In a network model with asymmetric recurrent connections it does not suffice to select the preferred connectivity direction of target neurons for individual source neurons depending on their positions. We also need to define how exactly the ’directions’ (*ϕ*) are distributed in space. For this, we considered four qualitatively different configurations *Homogeneous configuration:* In this configuration all neurons had the same *ϕ*, indicating a single-direction bias of the projections of all neurons (Figure 1c, left).

#### Random configuration

In this configuration *ϕ* for each neuron was chosen independently at random from a uniform distribution (Figure 1c, middle).

#### Perlin configuration

In this configuration *ϕ* was also chosen from a uniform distribution as in the random configuration, but it was assigned to each neuron according to the Perlin noise - a class of gradient noise (Perlin 1985). The generation of Perlin noise is described below. This spatial distribution of *ϕ* ensured that neighboring neurons had similar preferred directions (Figure 1c, right). *Symmetric configuration:* In this configuration all neurons established connections in an isotropic manner, without any directional preference.

### Perlin noise

To generate Perlin noise we first created a *p* × *p* grid (Perlin grid) that covered the whole network (of size *N × N*; *N* = 100 I-networks and *N* = 120 for EI-networks). We defined 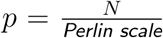. For example a *Perlin scale* = 20 meant that Perlin grid was of size 5 × 5 for the I-networks and 6 × 6 for the EI-networks. The variable *Perlin scale* controlled the spatial correlations. After defining the Perlin grid, each grid point was assigned a value chosen from a uniform distribution 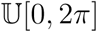. Next, we interpolated the Perlin grid to a size of the *N × N* (same size as the I-network or EI-network). The resulting values were used as the *ϕ* of the neuron located at that grid point. For more details about generation of Perlin noise please see (Perlin 1985).

### Input and network dynamics

All neurons received independent, homogeneous excitatory inputs from an external drive. We selected Gaussian white noise as an adequate input for generating ongoing spiking activity dynamics, which could be set to different activity levels by varying the input mean and variance independently.

### Identification of spatio-temporally clustered activity

To identify the STAS we rendered the spiking activity in a three dimensional space spanned by two spatial dimensions of the network and one time dimension. Each spike is a point and an STAS is a cluster in this 3-D space. We used the density-based spatial clustering algorithm of applications with noise (DBSCAN) (Ester et al. 1996) to determine individual clusters of the spiking activity in space and time. The DBSCAN algorithm required two parameters for the analysis: the maximum distance between two points in a cluster (*eps*) and the minimum number of points required to form a cluster (*minPts*). This algorithm needed a supervised control and an adequate value for *eps*, depending on the average spatial and -temporal distance (i.e. firing rate) between spikes of the neurons. For instance, when the average firing rate of the neurons was too high, then multiple STAS could be coalesced into a single STAS. On the other hand, when average firing rate of the neurons were small a single STAS could be dissociated into multiple small STAS. To avoid such problems, we reduced the temporal scale of spikes by a factor 20 and 3 for I-networks and EI-networks, respectively. Note that, this temporal rescaling was not used for the estimation of other properties of the network activity dynamics. The *eps* value was set to 3 and 2 for I-networks and EI-networks, respectively. Using the DBSCAN we identified STAS in successive, overlapping time windows of duration 1 sec (overlap duration 0.9 sec).

### Spatial arrangement of locally clustered activity

For each identified cluster we calculated the spatial centroids of activity bumps observed in successive time windows of 1 sec. The vectors composed of these successive centroids described the successive spatial coordinates of the bump activity and, hence, reveal the movement of the bump activity. Using these vectors we plotted the pathways of the moving activity bumps in the networks spatial map.

### Quantification of activity bump movement

Keeping track of direction changes in bump movement is an adequate measurement for the dynamics of sequential and non-sequential activities. In travelling waves, activity bumps typically move in a single direction, whereas in spatially fixed patterns, activity bumps alternate directions erratically over very short time scales. Between these two extrema, activity bump movements changed their direction slowly. The direction changing of bump movement is given by:

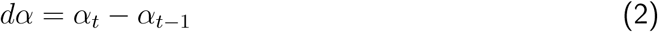

where *α_t_* denotes the direction of bump movement observed at time step *t*, and *dα* ranges from −π (opposite direction) over 0 (no alternation) to *π* (opposite direction).

### Identification of feedforward networks in the LCRN

To identify a feedforward network *FF* in the LCRN we started with a set of 64 neurons (*F_i_*), located in a 8 × 8 region in the network. This choice was motivated by seeing that individual spatial clusters of active neurons were of size 8 × 8. We then identified all post-synaptic neurons (*P_i_*) connected to any of the neurons in the set *F_i_*. From the set *P_i_* we selected 64 neurons (*F_i_*_+1_) that received the most number of connections from the *F_i_*. We repeated this procedure 50 times, starting at 100 different, randomly selected locations. Given the delays in the network, 50 time steps would imply that a sequence lasted for at least 100 ms. In this manner we identified feedforward networks with excitatory (inhibitory) connections from *F_n_* to *F_n_*_+1_ in EI-networks (I-networks).

To quantify the feedforward path we measured the number of neurons (*nFF*) belonging to *F*_1_… *F*_50_ over the trajectory between the centroids of *F*_1_ and *F*_50_ (Figure 7b,c). Note that each neuron was counted only once. The larger *nFF*, the longer and/or broader is the feedforward network.

In addition, we measured the *Effective length* as the Euclidean distance between the centroids of *F*_1_ and *F*_50_. Based on visual inspection of the locations of *F*_1_ … *F*_50_, we are implicitly assumed that {*F_n_*; *n* > 2} does not loop back to the same region where *F*_1_ is located.

To call the set of neurons that constitute *F*_1_ … *F*_50_ a feedforward path capable of creating STAS, we argued that {*F_n_*; *n* > 2} must be outside the connection region of the neurons in *F*_1_. In EI-networks the space constant of excitatory projections of a neurons is *σ_E_* = 12. If we assume that ≈ 70% connections are within one *σ_E_* (because the shape of connection probability function is Gaussian) then, in EI-networks the combined connection region of all neurons in *F*_1_ has a diameter of 12 + 8 + 12 = 32. Therefore, to be outside the connection region of *F*_1_, the centroid of *F*_50_ should be at least 16 grid points away from the centroid of *F*_1_ in EI-networks or the *Effective length* should be greater that 16. Similarly, we estimated the *Effective length* for I-networks as 16.

Thus, we defined that an effective feedforward pathway capable to creating STAS should have an *Effective length* > 16 (for EI- and I-networks). Finally, we defined *pFF* as the frequency of finding a feedforward path of *Effective length* > 16.

### Simulation Tools

All simulations of the network models were performed using the NEST simulation software (http://www.nest-initiative.org) (Peyser et al. 2017). The dynamical equations were integrated at a fixed temporal resolution of 0.1 ms. Simulation data were analyzed with Python using the scientific libraries SciPy (http://www.scipy.org) and NumPy (http://www.numpy.org), and visualized using the plotting library Matplotlib (http://matplotlib.org). The code will be made available at GitHub.

## Acknowledgements

We would like to thanks Dr. Upinder Singh Bhalla and Dr. Ulrich Egert for helpful discussions during the preparation of the manuscript. Partial funding of this work by the German-Isreali Foundation for Scientific Research and Development (GIF), the German Federal Ministry of Education and Research (BMBF) grant 01GQ0830 to the Bernstein Focus Neurotechnology (BFNT) Freiburg/Tuebingen, the Carl Zeiss Foundation, Parkinsonfonden Sweden, Swedish Research Council (StratNeuro and India-Sweden collaboration grant) is gratefully acknowledged.

## Supplementary materials

**Figure S1:**
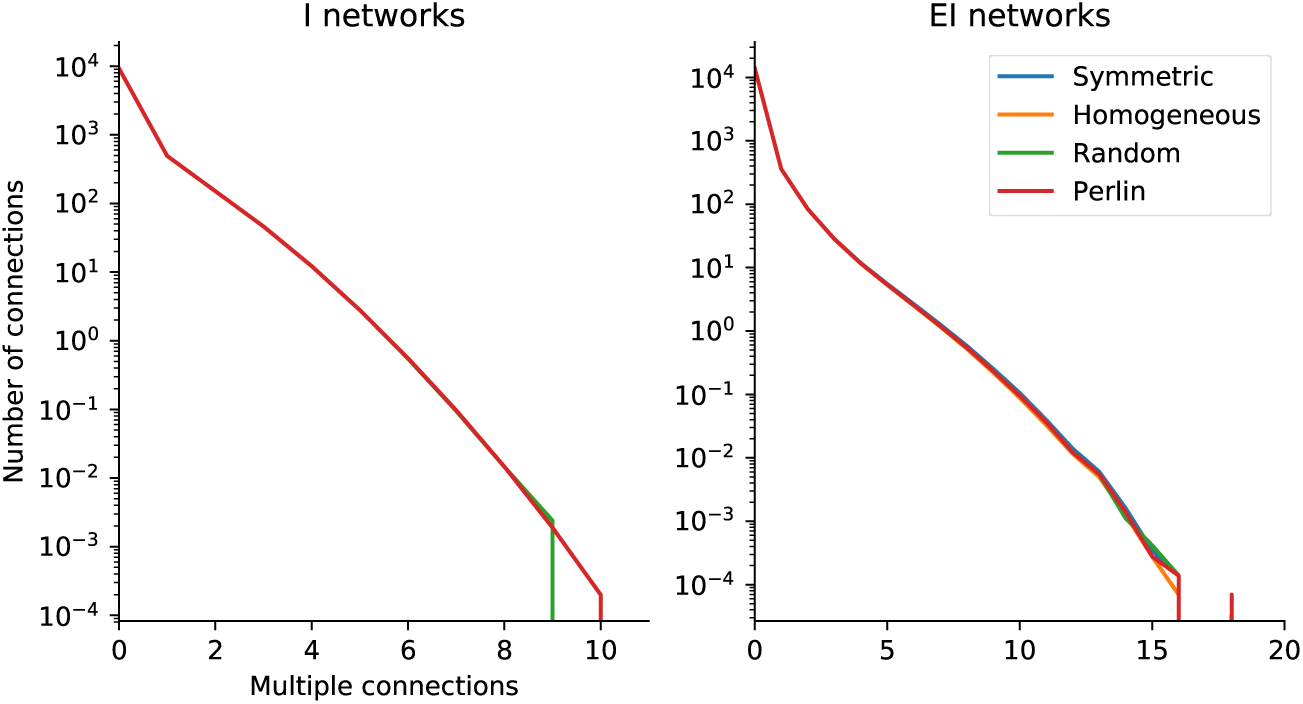
Multiplicity of connections. Count distribution of multiple connections between any pair of neurons in an I-network (left) and in an EI-network (right). The multiple connections were formed primarily because of the connectivity rule (local connectivity). Note that the network configuration (as indicated by different colors of the curves) had only a minute influence on the distribution of multiple connections.

**Figure S2:**
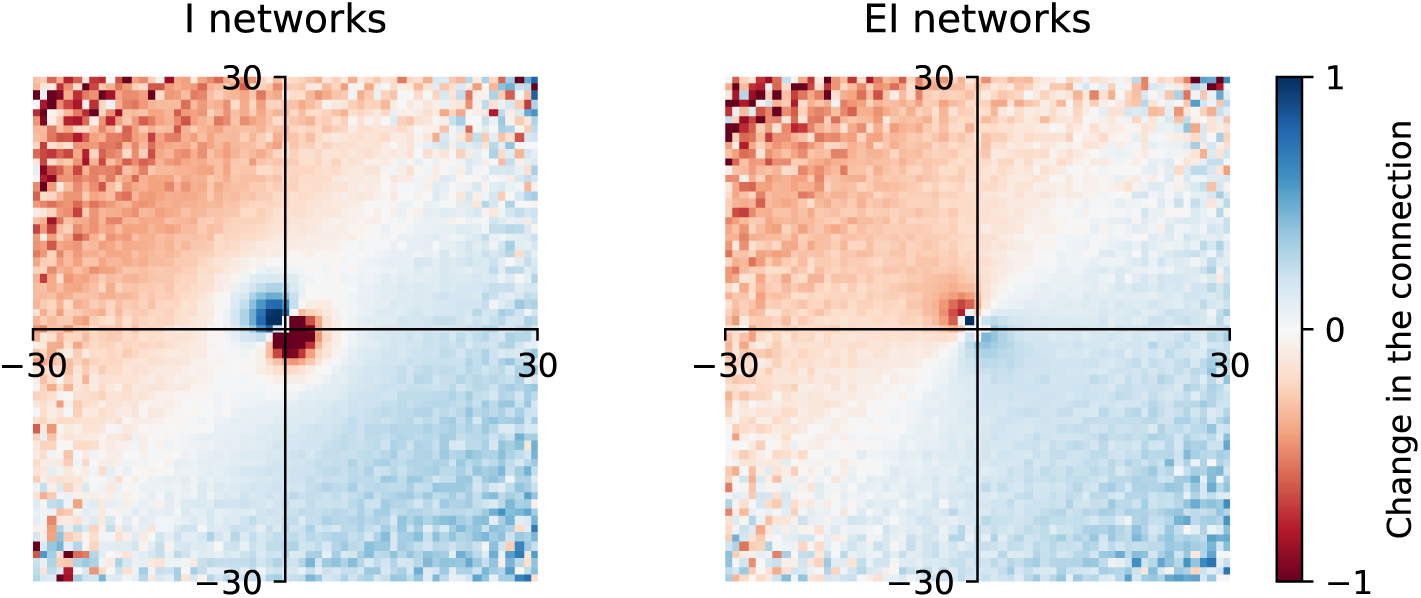
Effect of forcing a neuron to make some connections preferentially in the direction *ϕ*. To determine how the connectivity of a neuron changed when we forced it to make some connections preferentially in the direction *ϕ* while keeping its out-degree constant, we defined *δ_conn_ =* (*S − H)/S*, where *S* is the connectivity of a neuron in the symmetric configuration and *H* is the connectivity of the same neuron in the homogeneous configuration. That is, *δ_conn_* gives an estimate of the change in connectivity of a neuron relative to its connectivity in the homogeneous configuration. **Left** Average *δ_conn_* for I-networks (average over all the neurons in the network). **Right** Same as in the left panel, but for EI-networks. Forcing a neuron to make preferentially connections in the direction *ϕ* increased its connectivity in that direction by a factor 1 and, correspondingly, the connectivity was reduced by the same amount in the opposite direction. This change in the opposite direction is because we achieved asymmetry by shifting the connectivity cloud in the direction specified by *ϕ* (Figure 1). That is, in the immediate vicinity, the connection probability was doubled in the direction *ϕ*. This increase may look very large, but it nevertheless was not large enough to alter the probability of multiple connections in the network (Fig. S1). In the Figure we also notice a connectivity increase and corresponding decrease at larger distance as well, but this was not of much consequence because at such large distances the connection probability was very small to begin with.

**Figure S3:**
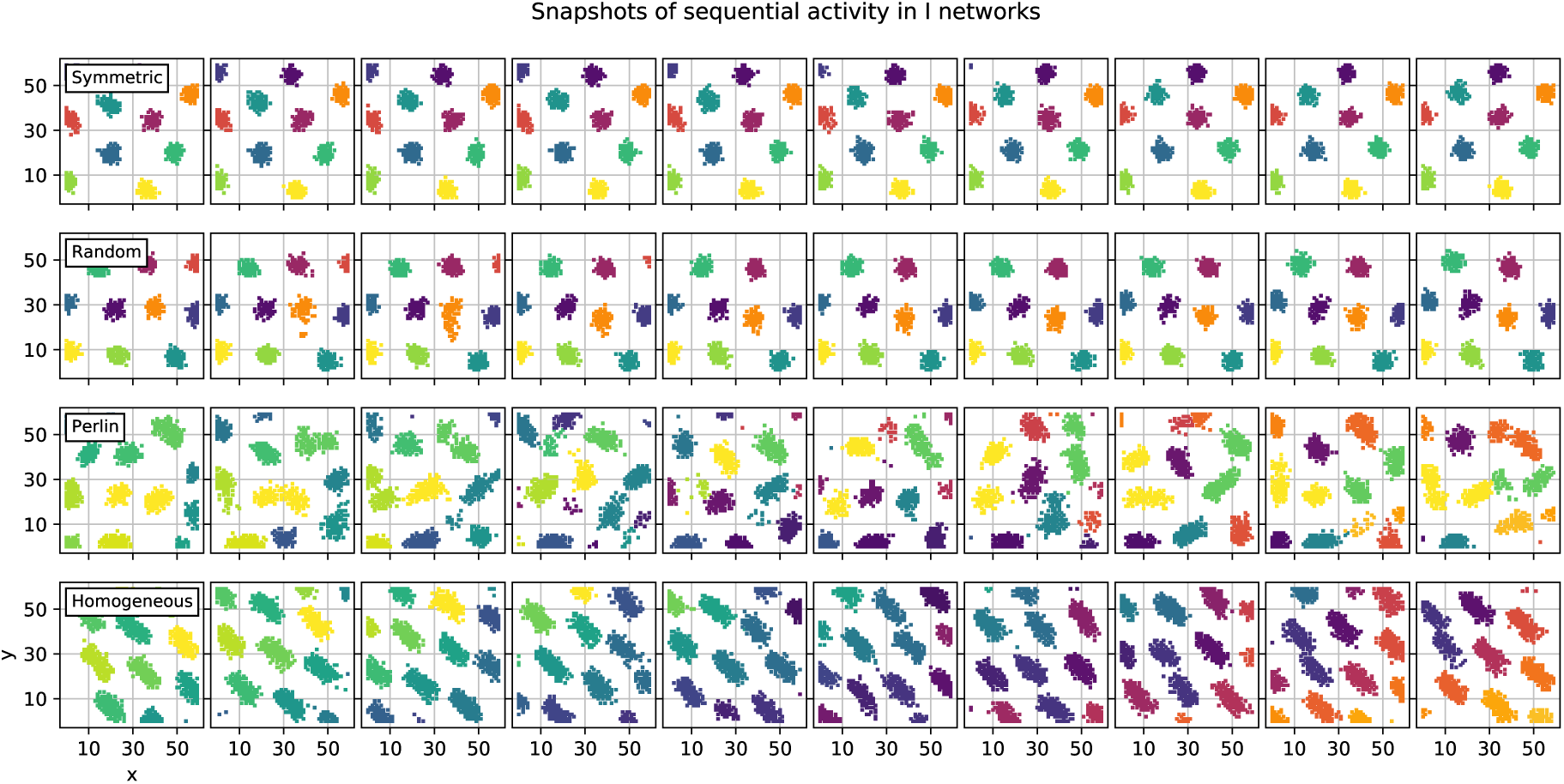
Snapshots of the spiking activity in I-networks for the four different configurations. Each row show the temporal evolution of the network activity for a different configuration. Each panel shows the activity of inhibitory neurons observed over a time window of 100 ms (disjoint windows), rendered in the two-dimensional network space. Each dot in a panel shows a spike of the neuron located at that grid point. Neurons are colored to identify individual spatio-temporal activity sequences (STAS). In each row, spikes rendered in the same color belong to the same STAS. Spikes that were not part of an STAS are not shown for clarity of presentation.

**Figure S4:**
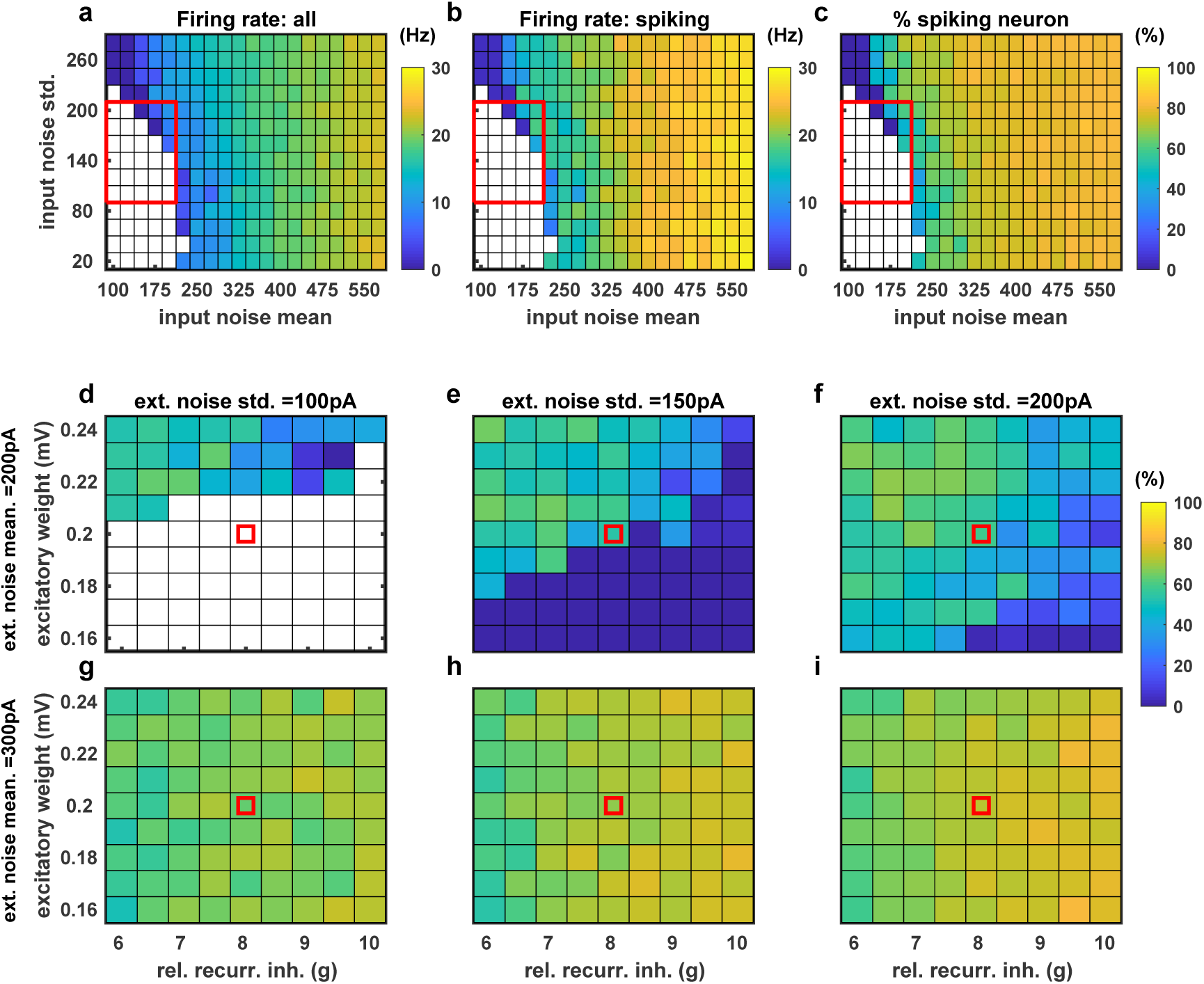
Effect of input and excitation-inhibition balance on the emergence of spatiotemporal sequences in an EI-network. **(a)** Average firing rate of all excitatory neurons as a function of the mean (x-axis) and standard deviation of the input noise to the neurons. **(b)** Same as in **a**, but here the average firing rates of only those excitatory neurons that spiked during an epoch of 2 sec is shown. **(c)** Fraction of excitatory neurons that spiked during an epoch of 2 sec. We used this fraction of spiking neuron as a proxy of STAS because, as shown in the main text, when networks were wired in the Perlin configuration, STASs emerged as soon as neurons crossed their spiking threshold. The value of EPSP amplitude and ’g’ used for panels **a-c** are marked by the red square in panels **d-i**. **(d)** Fraction of excitatory neurons that spiked as a function of the EPSP amplitude (y-axis) and ratio of the amplitudes of IPSP and EPSPs ’g’ (x-axis). Here we operated in an inhibition dominated regime. **(e-i)** Same as in panel **d**, but for different values of the input mean and standard deviation. The range of input mean and standard deviation used for panels **d-i** is marked by the red rectangle in panel **a**.

**Figure S5:**
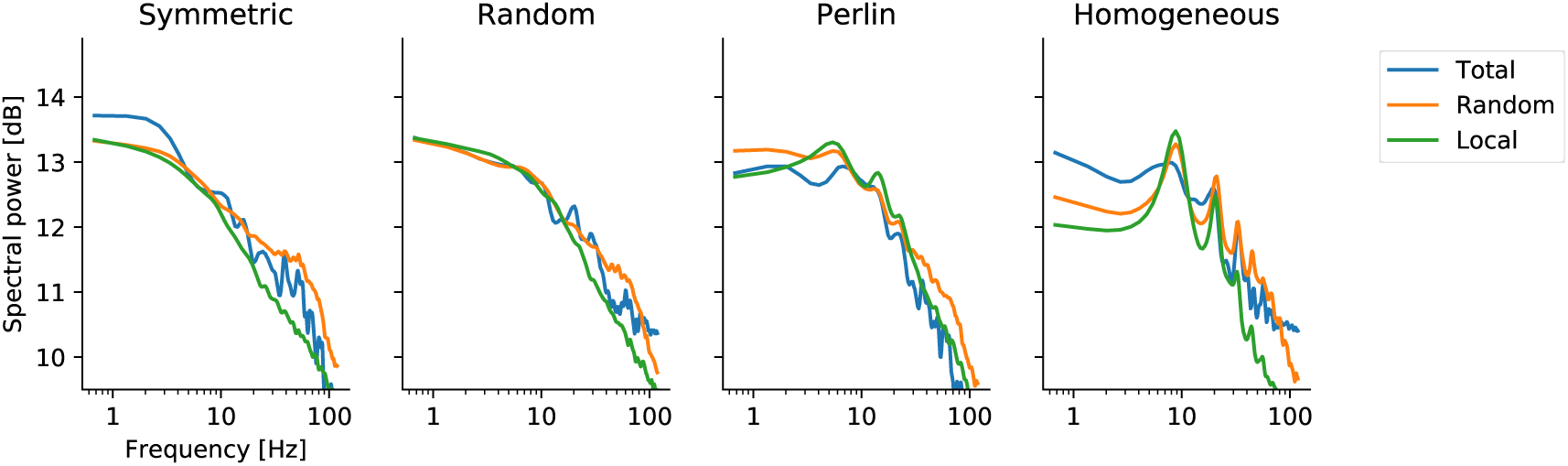
Power spectra of EI-network activity in different spatial inhomogeneity configurations. Power spectra of summed spiking activities of excitatory neurons (binwidth = 4ms), with different traces referring to the source of the data: the z-score of the spiking activity of the entire network population (blue trace), of 100 randomly selected neurons from the entire network (orange trace), and of the neurons located in a 10×10 region in the network (green trace). The spectral power in all network models peaked at approx. 60 Hz (gamma-band oscillations). Additionally, in network models with homogeneous and Perlin configurations, an additional, weak low-frequency peak, at around 4–6 Hz, appeared.

